## Supplemental_scripts for "CytofRUV: Removing unwanted variation to integrate multiple CyTOF datasets": CytofRUV_Figures.R

```
#title: "CytofRUV_Figures"
#author: "Marie Trussart"
#date: "31/03/2020"


library(CytofRUV)
library(CATALYST)
library(flowCore)
library(ggplot2)
library(readxl)
library(ruv)
library(purrr)
library(FlowSOM)
library(SummarizedExperiment)
library(ConsensusClusterPlus)
library(SingleCellExperiment)
library(shiny)
library(shinyjs)
library(shinydashboard)
library(writexl)


######################### Loading Metadata ######################### 
wd_data="/stornext/Bioinf/data/lab_speed/Marie/CytofRUV_Figures/"
metadata_filename="Metadata.xlsx"
panel_filename="Panel.xlsx"
seed=1234
clusters_nb=20
#Loading the data
data=CytofRUV::load_data(wd_data,metadata_filename,panel_filename)

## Cluster the data
set.seed(seed)
data$daf=cluster_data(data$daf,seed,markers_to_use=data$lineage_markers,clusters_nb)
saveRDS(data,paste(wd_data,"Raw_data_clustered.rds",sep=""))

## Raw data
raw_data <- data.frame(sample = data$daf$sample_id, cluster=cluster_ids(data$daf,"meta20"), t(SummarizedExperiment::assay(data$daf)))
saveRDS(raw_data,paste(wd_data,"Raw_Data.rds",sep=""))

## Proportions raw data
md=data$md
samples_order=md$sample_id[order(md$patient_id)]
raw_data$sample=factor(raw_data$sample,levels=samples_order)
counts_table <- table(raw_data$cluster, raw_data$sample)
props_table_raw <- t(t(counts_table) / colSums(counts_table)) * 100
saveRDS(props_table_raw ,paste(wd_data,"props_table_raw.rds",sep=""))

  
  
######################### ShinyApp on the raw data ######################### 
## Define parameters to use for R-Shiny
daf=data$daf
md=data$md
seed=1234
# Number of cells for diagnostic plots marker specific
n_subset_marker_specific <- 10000

# Define type of markers
daf_type <- daf[SingleCellExperiment::rowData(daf)$marker_class=="type", ]
daf_state <- daf[SingleCellExperiment::rowData(daf)$marker_class=="state", ]
sub_daf_state <- daf_state[, sample(ncol(daf_state), n_subset_marker_specific)]
sub_daf_type <- daf_type[, sample(ncol(daf_type), n_subset_marker_specific)]
# Define batch
batch_ids <- is.factor(rep(md$batch, nrow(daf)))
sampleID_sorted <- md$sample_id[order(md$patient_id)]

## Running Dimension Reduction -> TSNE
# Number of cells for tSNE plots marker specific
TSNE_subset <- 2000
print("Running TSNE")
set.seed(seed)
daf <- runDR(daf, "TSNE", cells = TSNE_subset)

# Number of cells for UMAP plots marker specific
UMAP_subset <- 2000
print("Running UMAP")
daf <- runDR(daf, "UMAP", cells = UMAP_subset)

## Launch Shiny
# For a subset of the data, define the number of cells for diagnostic plots
# n_subset <- 5000
# sub_daf <- daf[, sample(ncol(daf), n_subset)]

# For the full dataset: 
sub_daf <- daf
panel=data$panel

#CytofRUV::launch_Shiny()

  
  
######################### Figures before normalisation ######################### 
# MDS plot
library(ggrepel)
color_batch=c("#0072B2","#D55E00")

cs_by_s <- split(seq_len(ncol(daf)), daf$sample_id)
es <- as.matrix(assay(daf, "exprs"))
ms <- vapply(cs_by_s, function(cs) rowMedians(es[, cs, drop = FALSE]), 
        numeric(nrow(daf)))
rownames(ms) <- rownames(daf)
mds <- limma::plotMDS(ms, plot = FALSE)
df <- data.frame(MDS1 = mds$x, MDS2 = mds$y)
md <- metadata(daf)$experiment_info
m <- match(rownames(df), md$sample_id)
df <- data.frame(df, md[m, ])

png(paste(wd_data,"MDS_plot.png",sep=""))
    ggplot(df, aes_string(x = "MDS1", y = "MDS2", col = "batch")) + 
        geom_label_repel(aes_string(label = "sample_id",size=1.3), show.legend = FALSE) + 
        geom_point(alpha = 0.8, size = 1.5) + guides(col = guide_legend(overide.aes = list(alpha = 1, 
        size = 5))) + theme_void() + theme(aspect.ratio = 1, 
        panel.grid.major = element_line(color = "lightgrey", 
        size = 0.5),
        axis.title = element_text(face = "bold"), 
        axis.title.y = element_text(angle = 90), axis.text = element_text())+
        scale_color_manual(values = color_batch)+
        theme(text = element_text(size=20), legend.title = element_text(size = 20),
             legend.text = element_text(size = 20))
dev.off()

# Dendogram plot
png(paste(wd_data,"Dendogram_plot.png",sep=""))
plotExprHeatmap(daf, bin_anno = FALSE, row_anno = TRUE,)
dev.off()

# Markers distribution
png(paste(wd_data,"Lineage_Markers_distribution_plot_CLL2.png",sep=""))
    plotExprs(daf_type[, sample_ids(daf)%in%c("CLL2_B1","CLL2_B2")], color_by = "sample_id") +
      theme(axis.text=element_text(size=12),
            axis.title = element_text(size = 14),
            legend.title = element_text(size = 14),
            legend.text = element_text(size = 12)
      )
dev.off()

png(paste(wd_data,"Functional_Markers_distribution_plot_CLL2.png",sep=""))
    plotExprs(daf_state[, sample_ids(daf)%in%c("CLL2_B1","CLL2_B2")], color_by = "sample_id") +
      theme(axis.text=element_text(size=12),
            axis.title = element_text(size = 14),
            legend.title = element_text(size = 14),
            legend.text = element_text(size = 12)
      )
dev.off()

png(paste(wd_data,"Lineage_Markers_distribution_plot_HC1.png",sep=""))
    plotExprs(daf_type[, sample_ids(daf)%in%c("HC1_B1","HC1_B2")], color_by = "sample_id") +
      theme(axis.text=element_text(size=12),
            axis.title = element_text(size = 14),
            legend.title = element_text(size = 14),
            legend.text = element_text(size = 12)
      )
dev.off()

png(paste(wd_data,"Functional_Markers_distribution_plot_HC1.png",sep=""))
    plotExprs(daf_state[, sample_ids(daf)%in%c("HC1_B1","HC1_B2")], color_by = "sample_id") +
      theme(axis.text=element_text(size=12),
            axis.title = element_text(size = 14),
            legend.title = element_text(size = 14),
            legend.text = element_text(size = 12)
      )
dev.off()


## Heatmap clusters
png(paste(wd_data,"Heatmap_cluster_plot.png",sep=""))
plotClusterHeatmap(daf, hm2 = NULL, k = "meta20", m = NULL,
                       cluster_anno = TRUE, draw_freqs = TRUE)
dev.off()

## Tsne colored by batch for CLL1
png(paste(wd_data,"tsne_CLL1_colored_per_cluster_plot.png",sep=""))
plotDR(daf[, sample_ids(daf)%in%c("CLL1_B1","CLL1_B2")], "TSNE", color_by = "meta20") + facet_wrap("sample_id") +
      theme(axis.text=element_text(size=12),
            axis.title = element_text(size = 14),
            legend.title = element_text(size = 14),
            legend.text = element_text(size = 12))
dev.off()


# construct data.frame
xy <- reducedDim(daf, "TSNE")[,c(1, 2)]
colnames(xy) <- c("x", "y")
df <- data.frame(colData(daf), xy)
kids <- cluster_ids(daf, "meta20")
df[["meta20"]] <- kids
df <- df[!(is.na(df$x) | is.na(df$y)), ]

#### Plot tsne
dr <- data.frame(tSNE1 = df$x,tSNE2 = df$y)
dr$sample_id <-df$sample_id
dr$condition <- df$condition
dr$cell_clustering1 <- df$meta20
dr$batch=df$batch

## Plot CLL1 patient t-SNE colored per batch
#color_batch=c("#F8766D", "#00BFC4")
color_batch=c("#0072B2","#D55E00")
samples=c("CLL1_B1","CLL1_B2")
dr2=dr[dr$sample_id%in%samples,]
dr2$clusters=dr2$cell_clustering1
ggp <- ggplot(dr2,  aes(x = tSNE1, y = tSNE2, color = batch)) +
  geom_point(size = 0.8) +
  theme_bw() +
  scale_color_manual(values = color_batch) +
  guides(color = guide_legend(override.aes = list(size = 4), ncol = 2))
ggp + theme(aspect.ratio=1) +
  theme(text = element_text(size=20),
        axis.text.x = element_text(angle=90, hjust=1)) 
ggsave(paste(wd_data,"tsne-plot-samples_CLL1_colored_by_batch.png",sep=""))

## Plot CLL1 patient t-SNE colored by clusters
color_clusters <- c(
    "#DC050C", "#FB8072", "#1965B0", "#7BAFDE", "#882E72",
    "#B17BA6", "#FF7F00", "#FDB462", "#E7298A", "#E78AC3",
    "#33A02C", "#B2DF8A", "#55A1B1", "#8DD3C7", "#A6761D",
    "#E6AB02", "#7570B3", "#BEAED4", "#666666", "#999999",
    "#aa8282", "#d4b7b7", "#8600bf", "#ba5ce3", "#808000",
    "#aeae5c", "#1e90ff", "#00bfff", "#56ff0d", "#ffff00")
samples=c("CLL1_B1","CLL1_B2")
dr2=dr[dr$sample_id%in%samples,]
dr2$clusters=dr2$cell_clustering1
ggp <- ggplot(dr2,  aes(x = tSNE1, y = tSNE2, color = clusters)) +
  geom_point(size = 2.5) +
  theme_bw() +
  scale_color_manual(values = color_clusters,drop=FALSE) +
  guides(color = guide_legend(override.aes = list(size = 4), ncol = 2))
ggp + theme(aspect.ratio=1) +
  theme(text = element_text(size=20),
        axis.text.x = element_text(angle=90, hjust=1)) +
  facet_wrap(~ batch)+
  theme(aspect.ratio=1)
ggsave(paste(wd_data,"tsne-plot-samples_CLL1_colored_by_cluster_facet_batch.png",sep=""))

## Plot CLL1 patient t-SNE colored by clusters, batch1
dr3=dr2[dr2$batch=="1",]
ggp <- ggplot(dr3,  aes(x = tSNE1, y = tSNE2, color = clusters)) +
  geom_point(size = 2.5) +
  theme_bw() +
  scale_color_manual(values = color_clusters,drop=FALSE) +
  guides(color = guide_legend(override.aes = list(size = 4), ncol = 2))
ggp + theme(aspect.ratio=1) +
  theme(text = element_text(size=30),
        axis.text.x = element_text(angle=90, hjust=1)) +
  theme(aspect.ratio=1)+
  scale_x_continuous(limits=c(-30, 30))+
  scale_y_continuous(limits=c(-30, 30))
ggsave(paste(wd_data,"tsne-plot-samples_CLL1_colored_by_cluster_batch1.png",sep=""))

## Plot CLL1 patient t-SNE colored by clusters, batch2
dr3=dr2[dr2$batch=="2",]
ggp <- ggplot(dr3,  aes(x = tSNE1, y = tSNE2, color = clusters)) +
  geom_point(size = 2.5) +
  theme_bw() +
  scale_color_manual(values = color_clusters,drop=FALSE) +
  guides(color = guide_legend(override.aes = list(size = 4), ncol = 2))
ggp + theme(aspect.ratio=1) +
  theme(text = element_text(size=30),
        axis.text.x = element_text(angle=90, hjust=1)) +
  theme(aspect.ratio=1)+
  scale_x_continuous(limits=c(-30, 30))+
  scale_y_continuous(limits=c(-30, 30))
ggsave(paste(wd_data,"tsne-plot-samples_CLL1_colored_by_cluster_batch2.png",sep=""))

## Plot CLL1 patient t-SNE colored by batch, only cluster 9
dr4=dr2[dr2$clusters%in%c(9),]
ggp <- ggplot(dr4,  aes(x = tSNE1, y = tSNE2, color = batch)) +
  geom_point(size = 2.5) +
  theme_bw() +
  scale_color_manual(values = color_batch) +
  guides(color = guide_legend(override.aes = list(size = 4), ncol = 2))
ggp + theme(aspect.ratio=1) +
  theme(text = element_text(size=30),
        axis.text.x = element_text(angle=90, hjust=1)) +
  scale_x_continuous(limits=c(-30, 30))+
  scale_y_continuous(limits=c(-30, 30))
ggsave(paste(wd_data,"tsne-plot-samples_CLL1_cl9_colored_by_batch.png",sep=""))


## Abundances
png(paste(wd_data,"Proportions_per_cluster.png",sep=""))
samples_order=data$md$sample_id[order(data$md$patient_id)]
daf$sample_id=factor(daf$sample_id,levels=samples_order)
plotAbundances(daf, k = "meta20", by = "sample_id") +
      theme(axis.text=element_text(size=12),
            axis.title = element_text(size = 14),
            legend.title = element_text(size = 14),
            legend.text = element_text(size = 12),
            strip.text = element_blank()) +
      facet_wrap(facets = NULL, scales="fixed")
dev.off()

#### Plot per condition
props_table_raw=readRDS(paste(wd_data,"props_table_raw.rds",sep=""))
props <- as.data.frame.matrix(props_table_raw)
labels_col<-colnames(props)
dd=data.frame(cluster = rownames(props), props)
colnames(dd)=c("cluster",labels_col)
library(reshape2)
ggdf <- melt(dd,
             id.vars = "cluster", value.name = "proportion", variable.name = "sample_id")
ggdf$cluster <- factor(ggdf$cluster,levels=seq(1,20))
# Add condition and batch info
mm <- match(ggdf$sample_id, md$sample_id)
ggdf$condition <- factor(md$condition[mm])
ggdf$batch <- factor(md$batch[mm])

# Plot cluster proportions for all samples
ggdf2=ggdf[ggdf$cluster%in% c(2,6,7),]
ggdf2=ggdf2[ggdf2$sample_id%in%c("CLL1_B1","CLL1_B2"),]
png(paste(wd_data,"Barplot_cluster_proportions_CLL1_facet_cl2_6_7.png",sep=""))
ggplot(ggdf2, aes(x = batch, y = proportion, fill = cluster)) +
  geom_bar(stat = "identity") +
  facet_wrap(~cluster,scales = "free_y") +
  theme_bw() +
  theme(axis.text.x = element_text(angle = 90, hjust = 1)) +
  scale_fill_manual("cluster",values = color_clusters,drop=FALSE) +#+ theme(aspect.ratio=1) +
  theme(text = element_text(size=20))
dev.off()


######################### CytofRUV Normalisation CLL2_HC1_all_cl_20_k5 ######################### 
## Define parameters for normalisation
data=readRDS(paste(wd_data,"Raw_data_clustered.rds",sep=""))
daf=data$daf
md=data$md
dir_name_norm_data="CytofRUV_Norm_data_CLL2_HC1_all_cl_20_k5/"
raw_data=readRDS(paste(wd_data,"Raw_Data.rds",sep=""))
colnames(raw_data) <- gsub("^X", "",  colnames(raw_data))
rep_samples=list(c("CLL2_B1","CLL2_B2"),c("HC1_B1","HC1_B2"))
cluster_list_rep_samples <- list(seq(1,20),seq(1,20))
k_value <- 5
seed=1234

## CytofRUV normalisation
normalise_data(data=data,raw_data=raw_data,rep_samples=rep_samples, norm_clusters=cluster_list_rep_samples, k=k_value, num_clusters=clusters_nb,wd_data=wd_data,dir_norm_data=dir_name_norm_data)

## Define parameters to load and cluster the data
wd_norm=paste(wd_data,dir_name_norm_data,sep="")
metadata_norm_filename="Norm_Metadata.xlsx"
panel_norm_filename="Norm_Panel.xlsx"
seed=1234
clusters_nb=20

## Loading the norm data
norm_data=load_data(wd_norm,metadata_norm_filename,panel_norm_filename,cofactor=NULL)

## Cluster the norm data
set.seed(seed)
norm_data$daf=cluster_data(norm_data$daf,seed,markers_to_use=norm_data$lineage_markers,clusters_nb)
data <- data.frame(sample = norm_data$daf$sample_id, cluster=cluster_ids(norm_data$daf,"meta20"), t(SummarizedExperiment::assay(norm_data$daf)))
saveRDS(data,paste(wd_norm,"Norm_Data.rds",sep=""))

## Props table norm data
md=norm_data$md
samples_order=md$sample_id[order(md$patient_id)]
data$sample=factor(data$sample,levels=samples_order)
counts_table_norm <- table(data$cluster, data$sample)
props_table_norm <- t(t(counts_table_norm) / colSums(counts_table_norm)) * 100
saveRDS(props_table_norm ,paste(wd_norm,"props_table_norm.rds",sep=""))


######################### ShinyApp on the CytofRUV normalised data CLL2_HC1_all_cl_20_k5 ######################### 
daf=norm_data$daf
md=norm_data$md
daf_type <- daf[SingleCellExperiment::rowData(daf)$marker_class=="type", ]
daf_state <- daf[SingleCellExperiment::rowData(daf)$marker_class=="state", ]

# Number of cells for diagnostic plots marker specific
n_subset_marker_specific <- 10000
# Define batch
batch_ids <- is.factor(rep(md$batch, nrow(daf)))
sampleID_sorted <- md$sample_id[order(md$patient_id)]

## Running Dimension Reduction -> TSNE
# Number of cells for tSNE plots marker specific
TSNE_subset <- 2000
print("Running TSNE")
set.seed(seed)
daf <- runDR(daf, "TSNE", cells = TSNE_subset)

# Number of cells for UMAP plots marker specific
UMAP_subset <- 2000
print("Running UMAP")
daf <- runDR(daf, "UMAP", cells = UMAP_subset)

## Launch Shiny
# For a subset of the data, define the number of cells for diagnostic plots
# n_subset <- 5000
# sub_daf <- daf[, sample(ncol(daf), n_subset)]
  
# For the full dataset: 
sub_daf <- daf
panel=data$panel
  
#CytofRUV::launch_Shiny()


######################### Figures on the CytofRUV normalised data CLL2_HC1_all_cl_20_k5 ######################### 
# Markers distribution
png(paste(wd_norm,"Lineage_Markers_distribution_plot_CLL2.png",sep=""))
plotExprs(daf_type[, sample_ids(daf)%in%c("CLL2_B1","CLL2_B2")], color_by = "sample_id") +
  theme(axis.text=element_text(size=12),
        axis.title = element_text(size = 14),
        legend.title = element_text(size = 14),
        legend.text = element_text(size = 12)
  )
dev.off()

png(paste(wd_norm,"Functional_Markers_distribution_plot_CLL2.png",sep=""))
plotExprs(daf_state[, sample_ids(daf)%in%c("CLL2_B1","CLL2_B2")], color_by = "sample_id") +
  theme(axis.text=element_text(size=12),
        axis.title = element_text(size = 14),
        legend.title = element_text(size = 14),
        legend.text = element_text(size = 12)
  )
dev.off()

png(paste(wd_norm,"Lineage_Markers_distribution_plot_HC1.png",sep=""))
plotExprs(daf_type[, sample_ids(daf)%in%c("HC1_B1","HC1_B2")], color_by = "sample_id") +
  theme(axis.text=element_text(size=12),
        axis.title = element_text(size = 14),
        legend.title = element_text(size = 14),
        legend.text = element_text(size = 12)
  )
dev.off()

png(paste(wd_norm,"Functional_Markers_distribution_plot_HC1.png",sep=""))
plotExprs(daf_state[, sample_ids(daf)%in%c("HC1_B1","HC1_B2")], color_by = "sample_id") +
  theme(axis.text=element_text(size=12),
        axis.title = element_text(size = 14),
        legend.title = element_text(size = 14),
        legend.text = element_text(size = 12)
  )
dev.off()

## Tsne colored by batch for CLL1
png(paste(wd_norm,"tsne_CLL1_colored_per_cluster_plot.png",sep=""))
plotDR(daf[, sample_ids(daf)%in%c("CLL1_B1","CLL1_B2")], "TSNE", color_by = "meta20") + facet_wrap("sample_id") +
  theme(axis.text=element_text(size=12),
        axis.title = element_text(size = 14),
        legend.title = element_text(size = 14),
        legend.text = element_text(size = 12))
dev.off()

#### Plot tsne
# construct data.frame
xy <- reducedDim(daf, "TSNE")[,c(1, 2)]
colnames(xy) <- c("x", "y")
df <- data.frame(colData(daf), xy)
kids <- cluster_ids(daf, "meta20")
df[["meta20"]] <- kids
df <- df[!(is.na(df$x) | is.na(df$y)), ]

dr <- data.frame(tSNE1 = df$x,tSNE2 = df$y)
dr$sample_id <-df$sample_id
dr$condition <- df$condition
dr$cell_clustering1 <- df$meta20
dr$batch=df$batch

## Plot CLL1 patient t-SNE colored per batch
color_batch=c("#0072B2","#D55E00")
samples=c("CLL1_B1","CLL1_B2")
dr2=dr[dr$sample_id%in%samples,]
dr2$clusters=dr2$cell_clustering1
ggp <- ggplot(dr2,  aes(x = tSNE1, y = tSNE2, color = batch)) +
  geom_point(size = 0.8) +
  theme_bw() +
  scale_color_manual(values = color_batch) +
  guides(color = guide_legend(override.aes = list(size = 4), ncol = 2))
ggp + theme(aspect.ratio=1) +
  theme(text = element_text(size=20),
        axis.text.x = element_text(angle=90, hjust=1)) 
ggsave(paste(wd_norm,"tsne-plot-samples_CLL1_colored_by_batch.png",sep=""))

## Plot CLL1 patient t-SNE colored by clusters
color_clusters <- c(
  "#DC050C", "#FB8072", "#1965B0", "#7BAFDE", "#882E72",
  "#B17BA6", "#FF7F00", "#FDB462", "#E7298A", "#E78AC3",
  "#33A02C", "#B2DF8A", "#55A1B1", "#8DD3C7", "#A6761D",
  "#E6AB02", "#7570B3", "#BEAED4", "#666666", "#999999",
  "#aa8282", "#d4b7b7", "#8600bf", "#ba5ce3", "#808000",
  "#aeae5c", "#1e90ff", "#00bfff", "#56ff0d", "#ffff00")
samples=c("CLL1_B1","CLL1_B2")
dr2=dr[dr$sample_id%in%samples,]
dr2$clusters=dr2$cell_clustering1
ggp <- ggplot(dr2,  aes(x = tSNE1, y = tSNE2, color = clusters)) +
  geom_point(size = 0.8) +
  theme_bw() +
  scale_color_manual(values = color_clusters,drop=FALSE) +
  guides(color = guide_legend(override.aes = list(size = 4), ncol = 2))
ggp + theme(aspect.ratio=1) +
  theme(text = element_text(size=20),
        axis.text.x = element_text(angle=90, hjust=1)) +
  facet_wrap(~ batch)+
  theme(aspect.ratio=1)
ggsave(paste(wd_norm,"tsne-plot-samples_CLL1_colored_by_cluster_facet_batch.png",sep=""))

## Plot CLL1 patient t-SNE colored by clusters, batch1
dr3=dr2[dr2$batch=="1",]
ggp <- ggplot(dr3,  aes(x = tSNE1, y = tSNE2, color = clusters)) +
  geom_point(size = 0.8) +
  theme_bw() +
  scale_color_manual(values = color_clusters,drop=FALSE) +
  guides(color = guide_legend(override.aes = list(size = 4), ncol = 2))
ggp + theme(aspect.ratio=1) +
  theme(text = element_text(size=20),
        axis.text.x = element_text(angle=90, hjust=1)) +
  theme(aspect.ratio=1)+
  scale_x_continuous(limits=c(-30, 30))+
  scale_y_continuous(limits=c(-30, 30))
ggsave(paste(wd_norm,"tsne-plot-samples_CLL1_colored_by_cluster_batch1.png",sep=""))

## Plot CLL1 patient t-SNE colored by clusters, batch2
dr3=dr2[dr2$batch=="2",]
ggp <- ggplot(dr3,  aes(x = tSNE1, y = tSNE2, color = clusters)) +
  geom_point(size = 0.8) +
  theme_bw() +
  scale_color_manual(values = color_clusters,drop=FALSE) +
  guides(color = guide_legend(override.aes = list(size = 4), ncol = 2))
ggp + theme(aspect.ratio=1) +
  theme(text = element_text(size=20),
        axis.text.x = element_text(angle=90, hjust=1)) +
  theme(aspect.ratio=1)+
  scale_x_continuous(limits=c(-30, 30))+
  scale_y_continuous(limits=c(-30, 30))
ggsave(paste(wd_norm,"tsne-plot-samples_CLL1_colored_by_cluster_batch2.png",sep=""))

## Plot CLL1 patient t-SNE colored by batch, only cluster 9
dr4=dr2[dr2$clusters%in%c(17),]
ggp <- ggplot(dr4,  aes(x = tSNE1, y = tSNE2, color = batch)) +
  geom_point(size = 0.8) +
  theme_bw() +
  scale_color_manual(values = color_batch) +
  guides(color = guide_legend(override.aes = list(size = 4), ncol = 2))
ggp + theme(aspect.ratio=1) +
  theme(text = element_text(size=20),
        axis.text.x = element_text(angle=90, hjust=1)) +
  scale_x_continuous(limits=c(-30, 30))+
  scale_y_continuous(limits=c(-30, 30))
ggsave(paste(wd_norm,"tsne-plot-samples_CLL1_cl17_colored_by_batch.png",sep=""))


######################### Metrics of performance ######################### 
## Raw files
raw_files=paste(wd_data,"Raw_Data.rds",sep="")
props_table_raw=readRDS(paste(wd_data,"props_table_raw.rds",sep=""))
## Norm data k3
wd_norm=paste(wd_data,dir_name_norm_data,sep="")
props_table_norm=readRDS(paste(wd_norm,"props_table_norm.rds",sep=""))
## Define parameters and sample to compute the silhouette and LDA
nb_cells=10000
rep_samples=c("CLL2_B1","CLL2_B2")
## On specific cluster
cluster_to_select_raw_data=c(9,2)
cluster_to_select_norm_data=c(17,14)

############################ METRICS FOR EVALUATION OF METHODS ############################
Metrics_of_performance <- function(raw_files,cluster_to_select_raw_data,norm_files,outdir,cluster_to_select_norm_data,rep_samples,nb_cells=10000,props_table_raw,props_table_norm){
  ### METRICS TO ANALYSE PERFORMANCE OF THE NORMALISATION
  library(cluster)
  library(MASS)
  library(ruv)
  library(rsvd)
  library("gridExtra")
  library("ggpubr")
  library("tidyverse")
  library("RColorBrewer")
  library(purrr)
  library(matrixStats)
  library(ggplot2)
  library(reshape2)
  library(RColorBrewer)
  library(pheatmap)
  
  files <- c(raw_files, norm_files)
  file=c(1,2)
  data_list <- map(files, ~readRDS(.))
  raw_data=data_list[[1]]
  norm_data=data_list[[2]]
  colnames(norm_data) <- gsub("^X", "",  colnames(norm_data))
  colnames(raw_data) <- gsub("^X", "",  colnames(raw_data))
  nb_markers=dim(raw_data)[2]-2
  nclust=dim(props_table_norm)[1]
  nsample=dim(props_table_norm)[2]
  all_samples=colnames(props_table_norm)
  all_cluster=rownames(props_table_norm)
  
  ############################ 1.Silhouette and LDA on the Silhouette
  silhouette_specific_clusters <- function (cluster_to_select,rep_samples,nb_cells,data)
  {
    sample_ids=data$sample
    cell_clustering1=data$cluster
    expr= as.matrix(data[, 3:ncol(data)])
    colnames(expr) <- gsub("^X", "",  colnames(expr))
    rep_sample_ids=sample_ids[sample_ids%in%rep_samples]
    rep_expr=expr[sample_ids%in%rep_samples,]
    rep_cell_clustering=cell_clustering1[sample_ids%in%rep_samples]
    rep_sample_ids=rep_sample_ids[rep_cell_clustering%in%cluster_to_select]
    rep_expr=rep_expr[rep_cell_clustering%in%cluster_to_select,]
    rep_cell_clustering=rep_cell_clustering[rep_cell_clustering%in%cluster_to_select]
    ## Overall effect of bio vs batch with subsampling
    rep_data=data.frame(rep_expr,clust=as.numeric(rep_cell_clustering),batch=as.numeric(substr(rep_sample_ids,nchar(as.character(rep_sample_ids)), nchar(as.character(rep_sample_ids)))))
    colnames(rep_data) <- gsub("^X", "",  colnames(rep_data))
    base::set.seed(1234)
    subs_rep_data=rep_data[sample(nrow(rep_data), nb_cells), ]
    ## Silhouette
    over_ss_biology=silhouette(subs_rep_data$clust,dist(subs_rep_data[,1:ncol(data)]))
    over_ss_batch=silhouette(subs_rep_data$batch,dist(subs_rep_data[,1:ncol(data)]))
    over_ss<- c(bio = mean(over_ss_biology[,"sil_width"]), batch = mean(over_ss_batch[,"sil_width"]))
    #saveRDS(over_ss, paste(outdir,"Sil_Rep_",paste(c(rep_samples),collapse="_"),"_Clust_",paste(c(cluster_to_select),collapse="_"),"_",datatype,"_data_over_ss.rds",sep=""))
    return (over_ss)
  }
  
  silhouette_and_LDA_specific_clusters <- function (cluster_to_select,rep_samples,nb_cells,data,datatype){
    sample_ids=data$sample
    cell_clustering1=data$cluster
    expr= as.matrix(data[, 3:ncol(data)])
    colnames(expr) <- gsub("^X", "",  colnames(expr))
    rep_sample_ids=sample_ids[sample_ids%in%rep_samples]
    rep_expr=expr[sample_ids%in%rep_samples,]
    rep_cell_clustering=cell_clustering1[sample_ids%in%rep_samples]
    rep_sample_ids=rep_sample_ids[rep_cell_clustering%in%cluster_to_select]
    rep_expr=rep_expr[rep_cell_clustering%in%cluster_to_select,]
    rep_cell_clustering=rep_cell_clustering[rep_cell_clustering%in%cluster_to_select]
    ## Overall effect of bio vs batch with subsampling
    rep_data=data.frame(rep_expr,clust=as.numeric(rep_cell_clustering),batch=as.numeric(substr(rep_sample_ids,nchar(as.character(rep_sample_ids)), nchar(as.character(rep_sample_ids)))))
    colnames(rep_data) <- gsub("^X", "",  colnames(rep_data))
    base::set.seed(1234)
    subs_rep_data=rep_data[sample(nrow(rep_data), nb_cells), ]
    ## Silhouette
    over_ss_biology=silhouette(subs_rep_data$clust,dist(subs_rep_data[,1:ncol(data)]))
    over_ss_batch=silhouette(subs_rep_data$batch,dist(subs_rep_data[,1:ncol(data)]))
    over_ss<- c(bio = mean(over_ss_biology[,"sil_width"]), batch = mean(over_ss_batch[,"sil_width"]))
    saveRDS(over_ss, paste(outdir,"Sil_Rep_",paste(c(rep_samples),collapse="_"),"_Clust_",paste(c(cluster_to_select),collapse="_"),"_",datatype,"_data_over_ss.rds",sep=""))
    ### LDS plot
    batches=c(1,2)
    batch_data=subs_rep_data[subs_rep_data$batch%in%batches,]
    subs.lda2batch <- lda(clust + batch ~., data =batch_data )
    subs.lda.values <- predict(subs.lda2batch, batch_data[,1:ncol(data)])
    #convert to data frame
    color_batch=c("#0072B2","#D55E00")
    newbatch_data <- data.frame(batch = factor(batch_data$batch),cluster = factor(batch_data$clust), LDA1 = subs.lda.values$x[,1],LDA2 = subs.lda.values$x[,2],samp=paste("Rep_",batch_data$batch,"_Clust_",batch_data$clust,sep=""))
    ggplot(newbatch_data) + geom_point(aes(LDA1,LDA2,colour = batch,shape=cluster), size = 2.5) +
      scale_color_manual(values = color_batch) +
      guides(col = guide_legend(overide.aes = list(alpha = 1, size = 5))) +
      theme_bw() +
      theme(aspect.ratio = 1,
            panel.grid.minor = element_blank(),
            panel.grid.major = element_line(color = "lightgrey",size = 0.5))+
      theme(text = element_text(size=20), legend.title = element_text(size = 20),
            legend.text = element_text(size = 20),
            axis.text.x = element_text(angle=90, hjust=1))
    ggsave(paste(outdir,"LDA_Rep_",paste(c(rep_samples),collapse="_"),"_Clust_",paste(c(cluster_to_select),collapse="_"),"_",datatype,"_data.png",sep=""))
  }
  
  # ## All samples
  ind=1
  sil_raw_all_samples=matrix(nrow=nsample/2,ncol=3)
  colnames(sil_raw_all_samples)=c("samples","bio","batch")
  sil_raw_all_samples[,1]=substring(colnames(props_table_norm)[seq(1,nsample,2)],first=1,last = 4)
  sil_norm_all_samples=sil_raw_all_samples
  for (s in seq(1,nsample,2)){
    rep_samples_sel=colnames(props_table_norm)[c(s,s+1)]
    sraw=silhouette_specific_clusters(all_cluster,rep_samples_sel,nb_cells,data=data_list[[1]])
    sil_raw_all_samples[ind,2:3]=sraw
    snorm=silhouette_specific_clusters(all_cluster,rep_samples_sel,nb_cells,data=data_list[[2]])
    sil_norm_all_samples[ind,2:3]=snorm
    ind=ind+1
  }
  saveRDS(sil_raw_all_samples, paste(outdir,"Sil_all_samples_all_clust_raw_data_over_ss.rds",sep=""))
  saveRDS(sil_norm_all_samples, paste(outdir,"Sil_all_samples_all_clust_norm_data_over_ss.rds",sep=""))
  #
  
  ############################ 2. EMD
  EMD_metric <- function (data,binSize=0.005){
    nb_markers=dim(data[[1]])[2]
    distr <- list()
    for(file in c(1:length(files))){
      distr[[file]] <-  apply(data[[file]][,3:nb_markers],2,function(x){
        graphics::hist(x,
                       breaks = seq(-500,500,by=binSize),
                       plot = FALSE)$counts
      })
    }
    
    distances <- matrix(nrow=1,ncol=(nb_markers-3+1))
    colnames(distances)=colnames(data[[1]])[3:nb_markers]
    for(marker in c(1:(nb_markers-3+1))){
      distances[marker] <-
        emdist::emd2d(
          matrix(distr[[1]][,marker]),
          matrix(distr[[2]][,marker]))
    }
    return(distances)
  }
  
  ## EMD metric for rep samples per cluster
  EMD_per_cluster <- function(rep1,rep2){
    rep_comp_clust=matrix(nrow=nclust,ncol=nb_markers)
    colnames(rep_comp_clust)= colnames(data_list[[1]])[3:ncol(data_list[[1]])]
    row.names(rep_comp_clust)=paste("Cl_",seq(1,nclust),sep="")
    for (c in c(1:nclust)){
      rep1_clust_c=rep1[rep1$clust%in%c,]
      rep2_clust_c=rep2[rep2$clust%in%c,]
      rep_clust_c=list(rep1_clust_c,rep2_clust_c)
      rep_comp_clust[c,]=EMD_metric(rep_clust_c)
    }
    return(rep_comp_clust)
  }
  
  ## EMD metric for rep samples per cluster
  cluster_to_use=norm_data$cluster # !!!! IMPORTANT
  raw_data_tmp=raw_data
  raw_data_tmp$cluster=cluster_to_use
  raw_rep1=raw_data_tmp[raw_data$sample%in%rep_samples[1],]
  raw_rep2=raw_data_tmp[raw_data$sample%in%rep_samples[2],]
  norm_rep1=norm_data[norm_data$sample%in%rep_samples[1],]
  norm_rep2=norm_data[norm_data$sample%in%rep_samples[2],]
  raw_rep_comp_clust=EMD_per_cluster(raw_rep1,raw_rep2)
  norm_rep_comp_clust=EMD_per_cluster(norm_rep1,norm_rep2)
  saveRDS(raw_rep_comp_clust, paste(outdir,"EMD_metric_raw_data_per_cluster_",paste(c(rep_samples),collapse="_"),".rds",sep=""))
  saveRDS(norm_rep_comp_clust, paste(outdir,"EMD_metric_norm_data_per_cluster_",paste(c(rep_samples),collapse="_"),".rds",sep=""))
  ## EMD metric for all rep samples all cells
  ind=1
  EMD_metric_comp_raw_all_samples=matrix(nrow=nsample/2,ncol=nb_markers)
  colnames(EMD_metric_comp_raw_all_samples)=colnames(data_list[[1]])[3:ncol(data_list[[1]])]
  row.names(EMD_metric_comp_raw_all_samples)=substring(colnames(props_table_norm)[seq(1,nsample,2)],2)
  EMD_metric_comp_norm_all_samples=EMD_metric_comp_raw_all_samples
  for (s in seq(1,nsample,2)){
    rep_samples_sel=colnames(props_table_norm)[c(s,s+1)]
    raw_rep1=data_list[[1]][data_list[[1]]$sample%in%rep_samples_sel[1],]
    raw_rep2=data_list[[1]][data_list[[1]]$sample%in%rep_samples_sel[2],]
    norm_rep1=data_list[[2]][data_list[[1]]$sample%in%rep_samples_sel[1],]
    norm_rep2=data_list[[2]][data_list[[1]]$sample%in%rep_samples_sel[2],]
    raw_rep=list(raw_rep1,raw_rep2)
    norm_rep=list(norm_rep1,norm_rep2)
    EMD_metric_comp_raw_all_samples[ind,]=EMD_metric(raw_rep)
    EMD_metric_comp_norm_all_samples[ind,]=EMD_metric(norm_rep)
    ind=ind+1
  }
  saveRDS(EMD_metric_comp_raw_all_samples, paste(outdir,"EMD_metric_comp_raw_all_samples_all_cells.rds",sep=""))
  saveRDS(EMD_metric_comp_norm_all_samples, paste(outdir,"EMD_metric_comp_norm_all_samples_all_cells.rds",sep=""))
  EMD_all=rbind(EMD_metric_comp_raw_all_samples,EMD_metric_comp_norm_all_samples)
  ref=c(rep("raw",nsample/2),rep("norm",nsample/2))
  df=data.frame(EMD_all,ref,sample=rownames(EMD_all))
  ggdf <- melt(df, id.var = c("sample","ref"),
               value.name = "expression", variable.name = "antigen")
  #ggplot(ggdf,aes(x=expression,y=sample,color=ref))+geom_point(aes(colour = ref))+facet_wrap(~ref)
  ggplot(ggdf,aes(y=expression,x=sample,color=ref))+geom_boxplot(aes(colour = ref)) + theme(axis.text.x = element_text(angle = 45, hjust = 1))
  ggsave(paste(outdir,"EMD_metric_comp_raw_and_norm_all_samples_all_cells.png",sep=""))
  
  # ############################ 3. Clusters proportions accross rep
  # Compute different distances
  nclust=dim(props_table_norm)[1]
  nsample=dim(props_table_norm)[2]
  
  compute_dist <- function (props_table){
    nsample=dim(props_table)[2]
    nclust=dim(props_table)[1]
    Dist_props_table=matrix(nrow=5,ncol=nsample/2)
    colnames(Dist_props_table)=substring(colnames(props_table)[seq(1,nsample,2)],2)
    row.names(Dist_props_table)=c("Hell","TV","KL","BC","CW1")
    ind=1
    for (s in seq(1,nsample,2)){
      Dist_props_table[1,ind]=1/sqrt(2)*sqrt(sum((sqrt(props_table[,s])-sqrt(props_table[,s+1]))^2))
      Dist_props_table[2,ind]=1/2*sum(abs(props_table[,s]-props_table[,s+1]))
      Dist_props_table[3,ind]=sum(props_table[,s]*log(props_table[,s]/props_table[,s+1]))
      Dist_props_table[4,ind]=sum(sqrt(props_table[,s]*props_table[,s+1]))
      Dist_props_table[5,ind] = 1 - 4*sum(props_table[,s+1]/(props_table[,s]+props_table[,s+1])*
                                            (1-props_table[,s+1]/(props_table[,1]+props_table[,s+1])))/nclust
      ind=ind+1
    }
    return(Dist_props_table)
  }
  Dist_props_table_raw=compute_dist(props_table_raw)
  Dist_props_table_norm=compute_dist(props_table_norm)
  saveRDS(Dist_props_table_raw,paste(outdir,"Dist_props_table_raw_all_samples.rds",sep=""))
  saveRDS(Dist_props_table_norm,paste(outdir,"Dist_props_table_norm_all_samples.rds",sep=""))
}


Metrics_of_performance(raw_files,cluster_to_select_raw_data,paste(wd_norm,"Norm_Data.rds",sep=""),wd_norm,cluster_to_select_norm_data,rep_samples,nb_cells,props_table_raw,props_table_norm)
```
