## Supplemental_scripts for "CytofRUV: Removing unwanted variation to integrate multiple CyTOF datasets": CytofRUV_preprocessing_dataset.R

```
#title: "CytofRUV_preprocessing_dataset"
#author: "Marie Trussart"
#date: "31/03/2020"

library(readxl)
library(flowCore)
library(matrixStats)
library(ggplot2)
library(ggridges)
library(reshape2)
library(dplyr)
library(limma)
library(ggrepel)
library(FlowSOM)
library(cowplot)
library(lme4)
library(multcomp)
library(grid)
library(gridExtra)
library(CATALYST)

## Define work directory that contains the raw data
wd="CytofRUV_Figures_data/"
# Load the file containing all the fcs files 
file_list=read.table(paste(wd,"Raw_RUV1b_RUV3b/all_files.txt",sep=""),stringsAsFactors = F)
full_path_files=as.vector(paste(wd,"Raw_RUV1b_RUV3b/",file_list$V1,sep=""))
# Load all the files from the RUV1b run:
ff1 <- concatFCS(paste(wd,"RUV1b_Raw_data/",sep=""))
# Load all the files from the RUV3B run:
ff2 <- concatFCS(paste(wd,"RUV3B_Raw_data/",sep=""))
common_markers=intersect(colnames(ff1),colnames(ff2))
expr_ff2=exprs(ff2)
expr_ff2 = expr_ff2[, common_markers]
expr_ff1=exprs(ff1)
expr_ff1 = expr_ff1[, common_markers]
expr_ffall=rbind(expr_ff1,expr_ff2)
ffall=ff1
exprs(ffall)=expr_ffall
es <- flowCore::exprs(ffall)
es_t <- asinh(es/5)
chs <- flowCore::colnames(ffall)

####################### Functions adapted from Catalyst package ###########################
.get_ms_from_chs <- function(chs) {
  gsub("[[:punct:][:alpha:]]", "", chs)
}
ms <- .get_ms_from_chs(chs)
# find time, length, DNA and bead channels
time_col <- grep("time",        chs, ignore.case=TRUE)
lgth_col <- grep("length",      chs, ignore.case=TRUE)
dna_cols <- grep("Ir191|Ir193", chs, ignore.case=TRUE)

.get_bead_cols2 <- function(channels, beads) {
  ms <- gsub("[[:alpha:][:punct:]]", "", channels)
  if (is.character(beads)) {
    if (beads == "dvs") {
      bead_ms <- c(140, 151, 153, 165)#, 175)
    } else if (beads == "beta") {
      bead_ms <- c(139, 141, 159, 169, 175)
    }
  } else {
    bead_ms <- beads
  }
  n_beads <- length(bead_ms)
  bead_cols <- which(ms %in% bead_ms)
  if (length(bead_cols) != n_beads)
    stop("Not all bead channels found.")
  return(bead_cols)
}

.get_bead_inds <- function(x, y) {
  re <- assignPrelim(x, y, verbose=FALSE)
  re <- estCutoffs(re)
  re <- applyCutoffs(re)
  bc_ids(re) == "is_bead"
}

.update_bead_inds <- function(data, bead_inds, bead_cols, trim) {
  bead_es <- data[bead_inds, bead_cols]
  meds <- matrixStats::colMedians(bead_es)
  mads <- matrixStats::colMads(   bead_es) * trim
  ex <- matrixStats::colAnys(
    abs(t(bead_es) - meds) > mads)
  bead_inds[which(bead_inds)[ex]] <- FALSE
  bead_inds
}

.arrangeSmoothed <- function(x, p1, p2, out_path, fn, fn_sep, shiny=FALSE) {
    gt <- arrangeGrob(nrow=3, heights=c(5, .5, 5), widths=12,
        grobs=list(p1, rectGrob(gp=gpar(fill="white", col="white")), p2))
    if (shiny) {
        gt
    } else if (!is.null(out_path)) {
        if (is.null(fn)) {
            fn <- flowCore::description(x)$GUID
            fn <- gsub(".fcs", "", fn, TRUE)
        }
        out_nm <- paste(fn, "beads_before_vs_after.png", sep=fn_sep)
        png(file.path(out_path, out_nm), width=1500, height=1200, res=150)
        grid.draw(gt)
        dev.off()
    } else {
        grid.draw(gt)
    }
}

plotSmoothed <- function(df, main) {
  n <- length(chs <- colnames(df))
  baseline <- apply(df[, -1], 2, mean)
  p <- ggplot(df) + ggtitle(main) + theme_classic() +
    labs(x="Time [s]", y="Local median bead intensity") +
    theme(axis.text=element_text(size=10),
          plot.title=element_text(size=16, face="bold", hjust=.5),
          axis.title=element_text(size=12, face="bold"),
          panel.grid.major=element_line(color="grey"),
          panel.grid.minor=element_blank())

  for (i in seq_len(n)[-1])
    p <- p + geom_point(size=.5, alpha=.5,
                        aes_string(x=chs[1], y=chs[i], color=as.factor(chs[i]))) +
    geom_hline(show.legend=FALSE, lty=3, size=.5,
               aes_string(yintercept=baseline[i-1], col=as.factor(chs[i])))
  p + scale_color_manual(name=NULL,
                         values=RColorBrewer::brewer.pal(n-1, "Set1")) +
    guides(colour=guide_legend(override.aes=list(size=3)))
}

.arrangeSmoothed <- function(x, p1, p2, out_path, fn, fn_sep, shiny=FALSE) {
  gt <- arrangeGrob(nrow=3, heights=c(5, .5, 5), widths=12,
                    grobs=list(p1, rectGrob(gp=gpar(fill="white", col="white")), p2))
  if (shiny) {
    gt
  } else if (!is.null(out_path)) {
    if (is.null(fn)) {
      fn <- flowCore::description(x)$GUID
      fn <- gsub(".fcs", "", fn, TRUE)
    }
    out_nm <- paste(fn, "beads_before_vs_after.png", sep=fn_sep)
    png(file.path(out_path, out_nm), width=1500, height=1200, res=150)
    grid.draw(gt)
    dev.off()
  } else {
    grid.draw(gt)
  }
}


.outPlots <- function(x, es_t, bead_inds, remove, bead_cols, dna_cols,
                      smoothed_beads, smoothed_normed_beads, out_path, fn, fn_sep) {

  n <- nrow(es_t)
  n_beads <- length(bead_cols)

  p1 <- paste0(sprintf("%2.2f", sum(bead_inds)/n*100), "%")
  p2 <- paste0(sprintf("%2.2f", sum(remove)   /n*100), "%")
  t1 <- paste("Beads used for normalization (singlets only):", p1)
  t2 <- paste0("All events removed ",
               "(including bead-/cell-bead doublets): ", p2, "\n")
  ps <- c(
    .plotBeads(es_t, bead_inds, bead_cols, dna_cols, TRUE, FALSE, TRUE),
    .plotBeads(es_t, remove,    bead_cols, dna_cols, FALSE, TRUE, TRUE))

  if (!is.null(out_path)) {
    if (is.null(fn)) {
      fn <- flowCore::description(x)$GUID
      fn <- gsub(".fcs", "", fn, TRUE)
    }
    out_nm <- paste(fn, "beads_gated.pdf", sep=fn_sep)
    pdf(file.path(out_path, out_nm),
        width=n_beads*5, height=12.8)
    grid.arrange(grobs=ps, row=3, ncol=n_beads,
                 widths=rep(5, n_beads), heights=c(2.8, 5, 5),
                 top=textGrob(paste(t1, t2, sep="\n"), just="right",
                              gp=gpar(fontface="bold", fontsize=15)))
    dev.off()
  } else {
    grid.arrange(grobs=ps, row=3, ncol=n_beads,
                 widths=rep(5, n_beads), heights=c(3, 5, 5),
                 top=textGrob(paste(t1, t2, sep="\n"), just="right",
                              gp=gpar(fontface="bold", fontsize=15)))
  }
  p1 <- plotSmoothed(smoothed_beads, "Smoothed beads")
  p2 <- plotSmoothed(smoothed_normed_beads, "Smoothed normalized beads")
  .arrangeSmoothed(x, p1, p2, out_path, fn, fn_sep)
}

.outNormed <- function(ff, normed_es, remove_beads,
                       bead_inds, remove, out_path, fn, fn_sep) {
  if (is.null(fn)) {
    fn <- flowCore::description(ff)$GUID
    fn <- gsub(".fcs", "", fn, TRUE)
  }
  if (remove_beads) {
    ffs <- lapply(list(!remove, bead_inds, remove),
                  function(inds) new("flowFrame",
                                     exprs=normed_es[inds, ],
                                     parameters=flowCore::parameters(ff),
                                     description=flowCore::description(ff)))
    fs <- flowCore::flowSet(ffs)
    if (!is.null(out_path)) {
      if (is.null(fn)) {
        fn <- flowCore::description(ff)$GUID
        fn <- gsub(".fcs", "", out_nm, TRUE)
      }
      fn <- file.path(out_path, fn)
      out_nms <- paste0(c("normalised", "beads", "removed"), ".fcs")
      out_nms <- paste(fn, out_nms, sep=fn_sep)
      suppressWarnings(lapply(seq_len(3), function(i)
        flowCore::write.FCS(fs[[i]], out_nms[i])))
    }
    return(fs)
  } else {
    ff <- new("flowFrame",
              exprs=normed_es,
              parameters=flowCore::parameters(ff),
              description=flowCore::description(ff))
    if (is.null(out_path)) {
      return(ff)
    } else {
      out_nm <- paste0(fn, "normalised.fcs", sep=fn_sep)
      suppressWarnings(flowCore::write.FCS(ff, out_nm))
      return(ff)
    }
  }
}

.plotBeads <- function(es_t, bead_inds, bead_cols, dna_cols, hist, xlab, gate) {

    es <- sinh(es_t)*5
    chs <- colnames(es_t)
    n_beads <- length(bead_cols)

    df <- data.frame(
        es_t[, c(bead_cols, dna_cols)],
        id=as.numeric(bead_inds))
    if (gate) {
        gates <- data.frame(t(vapply(bead_cols, function(k)
            c(min(df[bead_inds, chs[k]]),
                max(df[bead_inds, chs[k]]),
                min(df[bead_inds, chs[dna_cols[1]]]),
                max(df[bead_inds, chs[dna_cols[1]]])),
            numeric(4))))
        colnames(gates) <- c("xmin", "xmax", "ymin", "ymax")
    }

    # sample 25'000 events for plotting
    N <- 25e3
    if (nrow(df) > N)
        df <- df[sample(nrow(df), N), ]

    # themes for plotting
    thms <- theme(
        aspect.ratio=1,
        panel.grid.major=element_line(color="grey"),
        panel.grid.minor=element_blank(),
        plot.margin=unit(c(0, .5, .5, .5), "cm"),
        axis.ticks.length=unit(.2, "cm"),
        axis.ticks=element_line(size=.5),
        axis.title=element_text(size=10),
        axis.text=element_text(size=8))

    # get axis limits
    x_max <- ceiling(max(es_t[, bead_cols])*2)/2
    y_max <- ceiling(max(es_t[, dna_cols ])*2)/2

    p <- list()
    if (hist) {
        for (i in seq_len(n_beads)) {
            med <- median(es[bead_inds, bead_cols[i]])
            min <- min(es[bead_inds, bead_cols[i]])
            max <- max(es[bead_inds, bead_cols[i]])
            med_t <- median(es_t[bead_inds, bead_cols[i]])
            min_t <- min(es_t[bead_inds, bead_cols[i]])
            max_t <- max(es_t[bead_inds, bead_cols[i]])
            p[[length(p)+1]] <- ggplot(data.frame(es_t[bead_inds, ])) +
                geom_histogram(aes_string(x=chs[bead_cols[i]], y="..ncount.."),
                    binwidth=.1, size=.025, fill="navy", col="cornflowerblue") +
                annotate("segment", min_t, 0, xend=min_t, yend=.1,  size=.75,
                    col="darkturquoise") +
                annotate("segment", med_t, 0, xend=med_t, yend=.05, size=.75,
                    col="darkturquoise") +
                annotate("segment", max_t, 0, xend=max_t, yend=.1,  size=.75,
                    col="darkturquoise") +
                annotate("text", .5, .75, hjust=0, size=3, fontface="italic",
                    col="darkslateblue", label=paste0(
                        "Median intensity: ", round(med, 2),
                        "\nLower boundary: ", round(min, 2),
                        "\nUpper boundary: ", round(max, 2))) +
                coord_cartesian(xlim=c(0, x_max), expand=FALSE) +
                scale_x_continuous(labels=function(x) format(x, nsmall=1)) +
                scale_y_continuous(breaks=c(0, .5, 1)) +
                ylab("Normalized counts") + theme_classic()  + thms +
                theme(axis.title.x=element_text(color="white"), aspect.ratio=.5)
            if (i != 1) p[[length(p)]] <- p[[length(p)]] +
                theme(axis.title.y=element_text(color="white"))
        }
    }
    for (i in seq_len(n_beads)) {
        p[[length(p)+1]] <- ggplot(df, aes_string(col="as.factor(id)",
            x=chs[bead_cols[i]], y=chs[dna_cols[1]])) + theme_bw() + thms +
            scale_colour_manual(values=c("0"="black", "1"="navy")) +
            geom_point(alpha=.25, size=1) + guides(color=FALSE) +
            coord_cartesian(xlim=c(0, x_max), ylim=c(0, y_max), expand=FALSE) +
            scale_x_continuous(labels=function(x) format(x, nsmall=1)) +
            scale_y_continuous(labels=function(x) format(x, nsmall=1)) +
            theme(panel.border=element_rect(colour="black", fill=NA, size=1))
        if (gate)
            p[[length(p)]] <- p[[length(p)]] + geom_rect(data=gates[i, ],
                aes_string(xmin="xmin", xmax="xmax", ymin="ymin", ymax="ymax"),
                col="darkturquoise", fill=NA, lty=3, size=1, inherit.aes=FALSE)
        if (!xlab) p[[length(p)]] <- p[[length(p)]] +
                theme(axis.title.x=element_text(color="white"))
        if (i != 1) p[[length(p)]] <- p[[length(p)]] +
                theme(axis.title.y=element_text(color="white"))
    }
    return(p)
}

.arrangeSmoothed <- function(x, p1, p2, out_path, fn, fn_sep, shiny=FALSE) {
  gt <- arrangeGrob(nrow=3, heights=c(5, .5, 5), widths=12,
                    grobs=list(p1, rectGrob(gp=gpar(fill="white", col="white")), p2))
  if (shiny) {
    gt
  } else if (!is.null(out_path)) {
    if (is.null(fn)) {
      fn <- flowCore::description(x)$GUID
      fn <- gsub(".fcs", "", fn, TRUE)
    }
    out_nm <- paste(fn, "beads_before_vs_after.png", sep=fn_sep)
    png(file.path(out_path, out_nm), width=1500, height=1200, res=150)
    grid.draw(gt)
    dev.off()
  } else {
    grid.draw(gt)
  }
}

.outPlots <- function(x, es_t, bead_inds, remove, bead_cols, dna_cols,
                      smoothed_beads, smoothed_normed_beads, out_path, fn, fn_sep) {

  n <- nrow(es_t)
  n_beads <- length(bead_cols)

  p1 <- paste0(sprintf("%2.2f", sum(bead_inds)/n*100), "%")
  p2 <- paste0(sprintf("%2.2f", sum(remove)   /n*100), "%")
  t1 <- paste("Beads used for normalization (singlets only):", p1)
  t2 <- paste0("All events removed ",
               "(including bead-/cell-bead doublets): ", p2, "\n")
  ps <- c(
    .plotBeads(es_t, bead_inds, bead_cols, dna_cols, TRUE, FALSE, TRUE),
    .plotBeads(es_t, remove,    bead_cols, dna_cols, FALSE, TRUE, TRUE))

  if (!is.null(out_path)) {
    if (is.null(fn)) {
      fn <- flowCore::description(x)$GUID
      fn <- gsub(".fcs", "", fn, TRUE)
    }
    out_nm <- paste(fn, "beads_gated.pdf", sep=fn_sep)
    pdf(file.path(out_path, out_nm),
        width=n_beads*5, height=12.8)
    grid.arrange(grobs=ps, row=3, ncol=n_beads,
                 widths=rep(5, n_beads), heights=c(2.8, 5, 5),
                 top=textGrob(paste(t1, t2, sep="\n"), just="right",
                              gp=gpar(fontface="bold", fontsize=15)))
    dev.off()
  } else {
    grid.arrange(grobs=ps, row=3, ncol=n_beads,
                 widths=rep(5, n_beads), heights=c(3, 5, 5),
                 top=textGrob(paste(t1, t2, sep="\n"), just="right",
                              gp=gpar(fontface="bold", fontsize=15)))
  }
  p1 <- plotSmoothed(smoothed_beads, "Smoothed beads")
  p2 <- plotSmoothed(smoothed_normed_beads, "Smoothed normalized beads")
  .arrangeSmoothed(x, p1, p2, out_path, fn, fn_sep)
}

.outNormed <- function(ff, normed_es, remove_beads,
                       bead_inds, remove, out_path, fn, fn_sep) {
  if (is.null(fn)) {
    fn <- flowCore::description(ff)$GUID
    fn <- gsub(".fcs", "", fn, TRUE)
  }
  if (remove_beads) {
    ffs <- lapply(list(!remove, bead_inds, remove),
                  function(inds) new("flowFrame",
                                     exprs=normed_es[inds, ],
                                     parameters=flowCore::parameters(ff),
                                     description=flowCore::description(ff)))
    fs <- flowCore::flowSet(ffs)
    if (!is.null(out_path)) {
      if (is.null(fn)) {
        fn <- flowCore::description(ff)$GUID
        fn <- gsub(".fcs", "", out_nm, TRUE)
      }
      fn <- file.path(out_path, fn)
      out_nms <- paste0(c("normalised", "beads", "removed"), ".fcs")
      out_nms <- paste(fn, out_nms, sep=fn_sep)
      suppressWarnings(lapply(seq_len(3), function(i)
        flowCore::write.FCS(fs[[i]], out_nms[i])))
    }
    return(fs)
  } else {
    ff <- new("flowFrame",
              exprs=normed_es,
              parameters=flowCore::parameters(ff),
              description=flowCore::description(ff))
    if (is.null(out_path)) {
      return(ff)
    } else {
      out_nm <- paste0(fn, "normalised.fcs", sep=fn_sep)
      suppressWarnings(flowCore::write.FCS(ff, out_nm))
      return(ff)
    }
  }
}

normCytof2 <- function(x, y,
                              out_path=NULL, fn=NULL, fn_sep="_",
                              remove_beads=TRUE, norm_to=NULL,
                              k=500, trim=5, verbose=TRUE, plot=TRUE) {

            # assure width of median window is odd
            if (k %% 2 == 0) k <- k + 1

            # check validity of 'out_path', 'fn', and 'fn_sep'
            stopifnot(is.null(out_path) ||
                        (is.character(out_path) & dir.exists(out_path)))
            stopifnot(is.null(fn) || is.character(fn))
            stopifnot(is.character(fn_sep))

            es <- flowCore::exprs(x)
            es_t <- asinh(es/5)
            chs <- flowCore::colnames(x)
            ms <- .get_ms_from_chs(chs)

            # find time, length, DNA and bead channels
            t_col <- grep("time", chs, ignore.case=TRUE)
            l_col <- grep("length", chs, ignore.case=TRUE)
            n_col <- grep("filenum", chs, ignore.case = TRUE)
            dna_cols <- grep("Ir191|Ir193", chs, ignore.case=TRUE)
            bead_cols <- .get_bead_cols2(chs, y)
            bead_chs <- chs[bead_cols]
            bead_ms <- ms[bead_cols]
            n_beads <- length(bead_ms)

            # identify bead singlets
            if (verbose) message("Identifying beads...")
            key <- data.frame(matrix(c(0, 0, rep(1, n_beads)), ncol=2+n_beads,
                                     dimnames=list("is_bead", c(191, 193, bead_ms))), check.names=FALSE)
            bead_inds <- .get_bead_inds(x, key)

            # get all events that should be removed later
            # including bead-bead and cell-bead doublets
            min_bead_ints <- matrixStats::colMins(es_t[bead_inds, bead_chs])
            remove <- apply(es_t[, bead_cols], 1, function(i) {
              above_min <- vapply(seq_len(n_beads),
                                  function(j) sum(i[j] > min_bead_ints[j]), numeric(1))
              sum(above_min) == n_beads
            })

            # trim tails
            bead_inds <- .update_bead_inds(es_t, bead_inds, bead_chs, trim)
            n_bead_events <- sum(bead_inds)

            bead_es <- es[bead_inds, bead_chs]
            bead_ts <- es[bead_inds, t_col]

            # get baseline (global mean)
            if (verbose) message("Computing normalization factors...")
            if (is.null(norm_to)) {
              baseline <- colMeans(bead_es)
            } else {
              if (is.character(norm_to)) {
                if (length(norm_to) != 1)
                  stop("'norm_to' should be a single character or flowFrame.")
                if (sum(flowCore::isFCSfile(norm_to)) != 1)
                  stop(norm_to, " is not a valid FCS file.")
                norm_to <- flowCore::read.FCS(norm_to,
                                              transformation=FALSE, truncate_max_range=FALSE)
              }
              chs_ref <- flowCore::colnames(norm_to)
              bead_cols_ref <- .get_bead_cols(chs_ref, y)
              bead_chs_ref <- chs_ref[bead_cols_ref]
              t_col_ref <- grep("time", chs_ref, ignore.case=TRUE)
              es_ref <- flowCore::exprs(norm_to)
              baseline <- colMeans(es_ref[, bead_chs_ref])
            }

            # smooth bead intensitites by conversion to local medians
            smoothed_beads <- apply(bead_es, 2, runmed, k, "constant")

            # compute slopes (baseline versus smoothed bead intensitites)
            # & linearly interpolate slopes at non-bead events
            bead_slopes <- rowSums(t(t(smoothed_beads)*baseline))/
              rowSums(smoothed_beads^2)
            slopes <- approx(bead_ts, bead_slopes, es[, t_col], rule = 2)$y

            # normalize raw bead intensities via multiplication with slopes
            normed_es <- cbind(
              es[,  c(t_col, l_col, n_col)],
              es[, -c(t_col, l_col, n_col)]*slopes)

            # smooth normalized beads
            bead_es_normed <- normed_es[bead_inds, bead_cols]
            smoothed_normed_beads <- apply(bead_es_normed, 2, runmed, k, "constant")

            # add time column
            smoothed_beads <- data.frame(bead_ts, smoothed_beads)
            smoothed_normed_beads <- data.frame(bead_ts, smoothed_normed_beads)
            colnames(smoothed_beads)[1] <-
              colnames(smoothed_normed_beads)[1] <- chs[t_col]

            if (plot)
              .outPlots(x, es_t, bead_inds, remove, bead_cols, dna_cols,
                        smoothed_beads, smoothed_normed_beads, out_path, fn, fn_sep)
            .outNormed(x, normed_es, remove_beads,
                       bead_inds, remove, out_path, fn, fn_sep)
          }

### Normalise the data using Finck normalisation on both runs together
norm_ffall=normCytof2(x=ffall, y="dvs", k=80,remove=F,out_path = paste(wd,"Norm_Raw_RUV1b_RUV3b_simult/",sep=""))
norm_ffall_without_beads=normCytof2(x=ffall, y="dvs", k=80,out_path = NULL,remove=T)
to_be_removed=norm_ffall_without_beads[[3]]
exp_toberem=exprs(to_be_removed)

### Debarcode Run1
norm_ff1=norm_ffall[1:dim(ff1)[1],]
exp_norm_ff1=exprs(norm_ff1)
exp_norm_ff1_without_beads=anti_join(as.data.frame(exp_norm_ff1),as.data.frame(exp_toberem))
exprs(norm_ff1)=as.matrix(exp_norm_ff1_without_beads)
ind=dim(ff1)[1]+1
data(sample_key)
re0 <- assignPrelim(x=norm_ff1, y=sample_key, verbose=TRUE)
re0
# estimate separation cutoffs
re <- estCutoffs(x=re0)
# use global separation cutoff
applyCutoffs(x=re, sep_cutoffs=0.35)
outFCS(x=re, y=norm_ff1,out_path=paste(wd,"Norm_Raw_RUV1b_RUV3b_simult/RUV1b/",sep=""))


### Debarcode Run2
norm_ff2=norm_ffall[ind:dim(norm_ffall)[1],]
exp_norm_ff2=exprs(norm_ff2)
exp_norm_ff2_without_beads=anti_join(as.data.frame(exp_norm_ff2),as.data.frame(exp_toberem))
exprs(norm_ff2)=as.matrix(exp_norm_ff2_without_beads)
data(sample_key)
re0 <- assignPrelim(x=norm_ff2, y=sample_key, verbose=TRUE)
re0
# estimate separation cutoffs
re <- estCutoffs(x=re0)
# use global separation cutoff
applyCutoffs(x=re, sep_cutoffs=0.35)
outFCS(x=re, y=norm_ff2,out_path=paste(wd,"Norm_Raw_RUV1b_RUV3b_simult/RUV3b/",sep=""))
```
