## Supplemental_scripts for "CytofRUV: Removing unwanted variation to integrate multiple CyTOF datasets": README.txt

This folder contains the scripts to generate the CytofRUV figures from the dataset FR-FCM-Z2L2. 

CytofRUV_preprocessing_dataset.R is the script that was used to generate the preprocessing on the dataset: data concatenation, normalisation and debarcoding are done using the R Catalyst package merging the two batches (RUV_1B and RUV3B) when applying the Finck normalisation. 

CytofRUV_Figures.R was used to generate the figures described in this manuscript and uses  
the CytofRUV R package with the Metadata.xlsx and Panel.xlsx files. The package is available at: www.github.com/mtrussart/CytofRUV. Installation and R code usage instructions for both the R package and the R-Shiny application can be found on the GitHub page. 
